# ICEPIC: A Toolkit to Discover Ice Binding Proteins from Sequence

**DOI:** 10.1101/2025.08.08.669420

**Authors:** Jimmy Zhang, Subbulakshmi Suresh, Shmuel Gleizer, Sophia Ewens, Aarya Venkat, Valentin Zulkower, Thomas Biernacki, Daniel Wen, Catherine Li, Mohammed Eslami, Susan Buckhout-White

**Affiliations:** Netrias, LLC., 1125 West Street, Suite 581, Annapolis MD 21401; Ginkgo Bioworks, Inc., 27 Drydock Ave 8th floor, Boston, MA 02210

**Keywords:** Ice binding proteins, antifreeze proteins, ice nucleation proteins, protein engineering

## Abstract

Ice binding proteins, such as antifreeze proteins (AFPs) and ice nucleation proteins (INPs), are critical for survival in subzero environments and have wide-ranging applications in biotechnology, agriculture, and materials science. Current discovery methods for these proteins are constrained by low throughput and limited datasets that are not conducive for engineering. Here, we present a high-throughput, sequence-based model that leverages contextual embeddings from protein language models to predict ice binding potential, as well as the expression and activity potential of candidate proteins. Using a curated data corpus of over 18,000 ice binding proteins — far larger than previous datasets — we fine-tuned a ProtBERT-based model, achieving 99% accuracy for prediction of ice binding potential. Sensitivity analyses through targeted mutagenesis (alanine and threonine substitutions) confirmed the model’s biological significance, revealing functionally important residues and sequence patterns. Additionally, we developed an expression prediction model that achieved an R^2^ score of 0.64 and low false-negative rates in identifying highly expressible candidates in *Pichia pastoris*. An additional regression model trained to predict ice activity as measured by thermal hysteresis achieved an R^2^ score of at least 0.79 with a clear difference in prediction between ice binding and non-ice binding proteins. Our toolkit advances the predictive accuracy, interpretability, and scalability of ice binding protein discovery, offering a powerful tool for protein engineering in cold-environment applications.

**Significance Statement:** Ice binding proteins enable organisms to survive freezing temperatures and are essential for applications in cryopreservation, agriculture, and materials science. However, discovering and engineering these proteins has been limited by small datasets and inadequate predictive tools. We developed a machine learning model trained on over 18,000 ice binding protein sequences to predict not only ice binding potential but also protein activity and expression in engineered hosts. This approach integrates advanced protein language models with biological context, enabling faster, more reliable discovery of ice binding proteins. Our platform advances rational protein design for real-world applications in climate resilience, biotechnology, and beyond.

## Introduction

Ice binding proteins, such as antifreeze proteins (AFPs), play a pivotal role in enabling organisms to survive in subzero environments by inhibiting ice crystal growth and modulating ice nucleation (1). Their relevance spans cryobiology, agriculture, food preservation, and synthetic material design. The ability to engineer these proteins carries important implications in cold-weather military, agricultural, shipping, and other industrial applications. To date, the identification of potential candidates has been achieved through statistical analyses with tools like BLAST and PSI-BLAST to identify similar sourced proteins (2). However, these methods often struggled due to the low sequence similarity among AFPs, which limited their predictive accuracy. Computational approaches can significantly enhance and accelerate the discovery of such proteins.

Recent computational approaches have significantly advanced the prediction of ice binding proteins by leveraging machine learning (ML) techniques. Dhibar and Jana introduced a high-performing method using n-gram features combined with an XGBoost classifier, utilizing penta-mers as a feature vector for predicting AFPs (3). While their model achieved outstanding performance, the use of a fixed-length motif (penta-mers) limits its capability to predict longer or non-contiguous dependencies. Usman *et al*. addressed the challenge of capturing non-linear sequence relationships through the use of latent space encoding of k-spaced amino acid pairs and a deep auto-encoder to extract significant features from the encoding scheme (4). However, the use of auto-encoders and specific hyperparameter tuning may lead to difficulty in broader application. Building on these foundations, Ali *et al*. developed an evolutionary-informed consensus model that integrates sequence motifs and evolutionary features, such as position-specific scoring matrices (PSSMs) (5). While this approach demonstrated robustness even on low-similarity datasets, its dependence on accurate multiple sequence alignments in PSSM generation could introduce bottlenecks, particularly for newly discovered or poorly characterized proteins. Complementing these strategies, Qi *et al*. proposed an ensemble model that integrates embeddings from multiple pre-trained transformer-based protein language models (ProtT5, ESM-1b, and Tape), further boosting predictive performance through an ensemble learning and soft voting mechanism (6). This model incorporates evolutionary relationships and implicit structural features from the pre-trained models. However, ensembling three large embedding models inflates inference time and GPU/CPU memory requirements, which may be impractical for high-throughput screens or lightweight deployments. Additionally, their study optimizes for a narrower end-point (binary AFP classification) rather than broader objectives such as relevance, expression level, or activity, which limits the applications for the model.

Collectively, these methods represent the state-of-the-art in AFP classification, highlighting a shift toward embedding-rich, interpretable, and evolution-aware modeling strategies (7). However, each of these models carry their own limitations. Many rely on fixed, handcrafted features that may not capture broader and deeper semantic relationships in protein sequences. Few models go beyond binary classification to predict expression likelihood or functional activity, which are critical for downstream applications. Finally, the training datasets for these models contain only a small number of Ice binding proteins, around 400-500, which further limits the utility and generalizability of these models. The models primarily focus on AFPs, but other proteins with ice binding properties exist, such as ice nucleation proteins (INPs). INPs are proteins that induce ice nucleation at temperatures above normal freezing point.

To address these gaps, we present a sequence-based language model that leverages contextual embeddings to predict not only ice binding features, but also the potential expression and activity of candidate proteins. This model is able to predict if a certain protein will interact with ice (*i*.*e*., ice growth inhibition or ice nucleation) based on the protein’s primary structure with a high level of confidence. This approach uses recent advances in protein language modeling to capture both global and local sequence dependencies, allowing for a more nuanced and generalizable understanding of ice protein-like behavior. Our dataset consists of over 18,000 ice binding proteins, capturing a wider spectrum of ice binding features. Overall, our model contributes to the growing toolkit for ice binding protein prediction and provides an interpretable, multi-faceted view of ice binding potential. With this toolkit, a protein engineering team can predict the likelihood of a *de novo* protein containing ice binding potential as well as its ice binding activity prior to testing. The toolkit can also be tailored to predict the expression from a specific host, the methylotrophic yeast *Pichia pastoris*. This toolkit can significantly streamline the downselection when engineering ice binding protein candidates. Thus, the model reported here will revolutionize the understanding of the structure-activity relationship of ice binding proteins and their subsequent formulation.

## Results

### Curation of a Data Corpus for Ice Binding Model

A significant challenge in training a sequence based model for specific ice binding functions is the sparsity of data available relevant to any specific function. Through metagenomic analyses, we realized that the challenge was less in the sparsity of data and more in the inconsistency of its annotations. A metagenomic screen was performed using 12 metagenomic seed proteins, which were curated to optimize for structural and taxonomic diversity based on literature. (please see ‘Metagenomic Library Curation’ in Methods for more information). An exploratory data analysis was then conducted to identify trends in both annotated names and functions of the proteins. These properties were obtained through their accession IDs in Uniprot and GenBank. The analysis identified a set of ice binding properties in their names and functions that were inconsistently annotated. Therefore, we developed a controlled vocabulary with distinct ice functions to harmonize all of these annotations. This included the following terms: *ice nucleation, ice-binding, ice structuring, and antifreeze* (**Supp. Figure A**). Note that these are general annotations conferred by the data in GenBank. Therefore, the GenBank ‘ice-binding’ annotation can be seen as a subset of the general category of ice binding proteins. We queried GenBank to curate a data corpus of all known annotated sequences that share these same properties in their annotations. In total, 18,573 protein sequences were obtained in this manner. All functions and names were harmonized to our controlled vocabulary (**Figure 2A**) (8). Along with the proteins in the 1st generation metagenomic library, this set of protein sequences was classified as ice binding protein sequences shown in **Figure 2B**. A similar procedure was used to curate an independent dataset from Swiss-prot, where the ice binding properties were used to query for proteins in Swiss-prot. After removing proteins already included in the ice binding sequences, there were a total of 37 proteins in this independent dataset. For sourcing the set of non-ice binding protein sequences shown in the same figure, a random set of proteins equal to the number of ice binding protein sequences was extracted from the proteomes of the 16 source organisms that were present in the metagenomic library. To eliminate potential data leakage between the two groups, any protein known to be a neighbor of an ice binding protein using data from StringDB or had an ice annotation was removed from the non-ice binding protein sequences. In total, this yields a corpus of over 39,000 proteins that are consistently annotated with a specific ice function. By comparison, similar datasets found in literature have approximately 400-500 AFPs with 7,000-9,000 non-AFPs (9, 4, 10). We provide this training corpus as supplementary data for use in sequence based modeling efforts.

### Prediction of Ice Binding Potential

Given a large data corpus, we can now fine tune a sequence based model to predict ice binding potential of a protein (**Figure 3A**). We wanted to first see if ice binding proteins, ones that have any of the above controlled vocabulary terms associated with them, had any distinct properties from non-ice binding proteins. A UMAP plot of the ProtBERT embeddings from the ice binding model data corpus shows a distinct pattern can be observed (**Figure 3B)** (11). As depicted by the two ovals in the figure, ProtBERT can discriminate between ice binding and non-ice binding clusters. The center of the UMAP plot depicts a prominent non-ice binding cluster (in the smaller oval), along with the great majority of metagenomic library proteins in the ice binding region surrounding the non-ice binding cluster in the larger oval. The center cluster in the smaller oval contains approximately 52% of the proteins, of which 88% of the cluster are non-ice binding and 1% are from the metagenomic library. Conversely, the region defined between the two ovals contains approximately 29% of the proteins, of which 6% of the region are non-ice binding and 84% are from the metagenomic library. This indicates that ProtBERT has already provided a significant distinction between the proteins in the metagenomic library and non-ice binding proteins. Fine-tuning layers added to ProtBERT will linearize this discrimination as shown by the UMAP after fine-tuning. The ice binding model achieved a 99% accuracy on a testing set, a 90% accuracy on a held-out dataset of the 1st generation metagenomic library, and an 89% accuracy on an independent dataset curated from Swiss-prot. This indicates the model is highly capable of discerning ice binding and non-ice binding proteins based on their sequences.

### Sensitivity Testing from Mutational Analysis of Proteins

Given the high performance of a sequence based model, the immediate question is whether the training data readily discriminates between ice/non-ice binding proteins. Given that non-ice binding proteins may have significantly different motifs from the ice binding proteins, the possibility of curating a dataset that is so obviously different that any ice binding model can discriminate between them is high. To answer this question, we wanted to test the sensitivity of the model to amino acid mutations. Most ice binding proteins rely on the amino acids threonine and/or alanine (T/A), where the sequence order, spacing, and composition of these amino acids aid in ordering water molecules, contributing to the formation of an ice lattice (9). Threonine-rich AFPs bind to ice through a combination of hydrogen bonding and hydrophobic interactions, facilitated by a specific arrangement of threonine residues and surrounding amino acids at the ice binding surface. They bind via anchored clathrate waters, positioning the water molecules similar to ice sheets (12, 13). According to literature, *Rm*AFP (*6XNR*), *Mp*AFP (*A1YIY3*), *Tm*AFP (*O16119*), and *Dc*AFP (*O46346*) are characterized as having threonine-dependent ice binding motifs (14, 12, 15). Meanwhile, *Gr*AFP (*A0A7U0TF39*) and *Sf*AFP (*Q38PT6*) are hypothesized to promote ice binding via an alanine dependent motif (16). For example, *Gr*AFP forms a polyproline type II helical bundle where alanine is the principal component of the ice binding surface. The small size of alanine’s methyl group has been shown to allow surface waters to interact with the peptide backbone, where crystallographic water molecules are observed in the troughs between alanine side chains.

To evaluate the functional role of threonine (T) and alanine (A) in selected ice-binding proteins, we performed targeted mutations in which each T or A residue was substituted with one of the 19 alternative amino acids. These “alanine-to-random” or “threonine-to-random” mutations involved replacing every occurrence of the specified amino acid with a randomly selected alternative. The effect of each mutation was quantified by the change in the model’s predicted ice-binding probability between the mutant and its corresponding original protein. This change is reported as the mean probability delta (or mean delta, as shown in the figures). Since the number of T and A residues varies across proteins, the number of mutations was normalized by the total number of T or A residues in the original sequence, resulting in a % mutation. For example, mutating all alanine residues in a protein corresponds to 100% alanine mutation. These mutations affected only a portion of each protein sequence. On average, alanine residues make up 13% of the total sequence length, while threonine residues comprise 12%.

As shown in **Figure 3C** (left column), the model responds to the mutated number percentage of alanine mutations through decreasing the mean delta (more negative). The label prediction changes (*i*.*e*., ice binding to non-ice binding) increased in a corresponding manner. As expected, the model has associated alanine as a crucial component of ice binding function, and mutating these alanines into another amino acid has resulted in a significant shift in the model’s predictions and label changes. Most of the proteins exhibit high sensitivity to the ‘alanine-to-random’ mutations, demonstrated by the greater amount of label changes and negative mean deltas. This is consistent with the hypothesis from literature that some of these proteins (*Gr*AFP and *Sf*AFP) have alanine-dependent ice binding motifs. Therefore, the mutation of alanine to another amino acid would theoretically decrease the ice binding potential of the protein. The exceptions are for *Mp*AFP and *Tm*AFP. For *Tm*AFP, the discrepancy may be due to the low ice binding model prediction, leading to a lower propensity for the label changes. For *Mp*AFP, the significantly greater size of the protein compared to other tested proteins may have been the cause, meaning that a lower percentage of this protein was mutated on average. The low significance of alanine for the ice binding function of these proteins may also be a factor. Based on literature, *Mp*AFP and *Tm*AFP are known to have threonine-dependent ice binding motifs, not alanine-dependent.

The model also displayed a response to the changing number of ‘threonine-to-random’ mutations (**Figure 3C**; right column). Similar to the ‘alanine-to-random’ mutations seen earlier, as the ‘threonine-to-random’ mutations increased, the mean probability deltas became more negative. The label changes increased correspondingly, similar to the trend seen with the ‘alanine-to-random’ mutations. However, a key difference between these types of mutations is the set of proteins that are most affected by the mutations. As noted in the ‘alanine-to-random’ mutational analysis, *Mp*AFP and *Tm*AFP are not sensitive to the alanine mutations. In contrast, these proteins display significantly higher sensitivity to the ‘threonine-to-random’ mutations, as seen in the increasingly negative mean probability deltas and increase in label changes compared to the rest of the proteins. This corroborates with current literature that these proteins have threonine-dependent ice binding planes (14, 12, 15). Overall, these results suggest that the ice binding model contains the biological context that is consistent with the literature.

We also used the ESM2 model to determine the effects of these mutations on ice binding proteins. Similar to ProtBERT, which is the base model for our workflow, ESM2 is a protein language model with BERT/transformer-based architecture (17). However, it was trained to determine the favorability of a mutation within an evolutionary context. A more favorable mutation generally indicates better protein stability and/or function. Overall, a comparison of the results between our ice binding model and ESM2 indicates that while similarities exist (*Sf*AFP from **Figure 3D**), there are also notable differences (*Tm*AFP from **Figure 3D**). For example, our model was able to determine that *Sf*AFP mutations from alanine to cysteine (C), phenylalanine (F), methionine (M), or tyrosine (Y) resulted in a major decrease in ice binding potential, which coincides with ESM2’s metrics for protein stability. On the other hand, our model determined that *Tm*AFP mutations from threonine to isoleucine (I), proline (P), or arginine (R) induced an increase in ice binding potential, which is contrary to ESM2 displaying a decrease in protein stability for the same mutations. An explanation for these discrepancies is the general objective for both models. ESM2’s objective was to build a zero-shot log odds ratio model where it compares whether or not a mutant will occur over a wild-type amino acid. The assumption of ESM2 is that mutations that are observed in nature are stable and functional. However, specific functions are ignored in ESM2. Instead, the log odds ratio is used as a proxy for favorability of whether or not the mutant is stable/functional. On the other hand, the ice binding model was trained to focus on a single ice binding objective, which assesses the likelihood a sequence is coming from an ice binding protein or not. It directly encodes the ice binding potential into the fine-tuning layers while ESM2 is a more generalized approach. This is shown in **Figure 3D** by the difference in the metrics’ scales. The ice binding metric is between 0 and 1 to indicate the probability of a sequence being ice binding. ESM2 utilizes a scoring method that follows a different scale based on the model’s evolutionary context. Objectively speaking, neither model is superior to the other, and both results should be analyzed in tandem for future implementation. However, this analysis highlighted a significant point from the results of the ice binding model: threonine-to-random mutations can increase the ice binding metric. While the ground truth cannot be verified at this moment, the general hypothesis was that removing threonine would decrease ice binding potential since it is known to be integral to ice function and is shown to decrease protein stability. It is possible that these mutants that were predicted to be more ice binding would not fold properly due to the lack of protein stability. Overall, when coupled with ESM2, the results suggest that different amino acids are more critical to the ice binding function of different proteins, and the model is able to successfully detect those differences.

### Prediction of Ice Binding Protein Expression from Sequence

While a significant portion of our work focused on the development of the sequenced-based ice binding model, the success of the ice binding model for engineering ice binding proteins could not be validated until they are effectively expressed in a host. The methylotrophic yeast *Pichia pastoris* was selected as the host system for protein expression due to several key advantages: its extensive use in literature, including for the production of ice-modulating proteins; its ability to achieve higher cell densities than *E. coli*; its capacity for extracellular protein secretion, which simplifies downstream purification; and its ability to perform post-translational modifications such as glycosylation, which has been implicated in ice-binding function (18, 19, 20, 21).

We have further enhanced *P. pastoris* expression by developing a proprietary platform that incorporates synthetic promoter systems, which significantly outperform existing state-of-the-art alternatives. Our proprietary platform also includes specialized base strains that are engineered to facilitate expression of proteins with disulfide bonds and to minimize protease activity.

Using our high-throughput strain engineering platform, single-copy integration of each construct was achieved and verified by sequencing. Protein expression was induced and quantified using a C-terminally fused HiBiT tag (Promega Corporation), an 11-amino-acid epitope that enables highly sensitive detection. Upon addition of LgBiT, the complementation partner, and substrate, a bioluminescent signal is produced, allowing relative expression levels to be measured. Strains are ranked based on this assay, and top candidates are advanced for fermentation in bioreactors.

In conjunction with our protein formulation efforts, we developed an ice expression model to predict the expression level via the HiBiT expression assay of the engineered proteins in the *P. pastoris* host. The expression model took as input the vector embeddings generated by ProtBERT to predict expression levels. A regression head was implemented, offering a baseline approach for learning the relationship between embeddings and expression. The expression model yielded an overall R^2^ at 0.64. We wanted to focus on the model predictions for highly-expressed protein candidates (expression value of over 0.25), as the main goal of this model is to accurately discern the actual highly-expressed candidate with a high expression prediction. Thus, we also isolated the highly-expressed candidates in our subsequent analysis and optimized the model based on this criterion. The model achieved an R^2^ for high expression candidates of 0.62. In a similar context, the false negative (FN) rate is the proportion at which actual highly-expressed proteins are predicted to have low-expression by our model. This rate was minimized so that the model can be as comprehensive as possible in capturing all possible highly-expressed protein candidates. The optimized model has an FN rate of 0.17. A prediction plot from this regression head model is shown in **Figure 4B**, which also highlights the optimizing criterion for the model.

A second metagenomic library that was not used in training was used to further validate the expression model. The correlation coefficients between the predicted and actual values are shown for each sub-library within the metagenomic library. Results, as shown in **Figure 4C**, indicate that the model was able to achieve a correlation coefficient of over 0.5 for 4 of the 6 sub-libraries, with INP sub-library displaying the best correlation at 0.75. Meanwhile, the AFP and P19614 sub-libraries display much lower correlations at 0.29 and 0.10, respectively. Representative scatter plots of predicted vs actual values for the INP and AFP sub-libraries are shown in **Figure 4C**, indicating the differences in correlation. The reason why the model struggled with predicting the expression values for the P19614 sub-library was due to the 1st generation metagenomic library lacking associated training context. A hypothesis for the low correlation for the AFP sub-library is that the huge diversity in sequence and structural motifs for AFP proteins is hard to capture for the model. A deeper analysis into the AFP sub-library was taken by grouping the library proteins by their associated AFP seed proteins and calculating their correlation coefficients similar to the methodology used earlier. The results for the deeper AFP analysis are shown in the right plot of **Figure 4C**. These results indicate noticeable failure points for the model, which includes the specific AFP groups of proteins that have negative correlation coefficients between predicted vs actual expression values. A closer analysis into these AFP groups and their associated AFP seeds may uncover additional information to further improve the model.

### Prediction of Ice Activity from Sequence

Due to the limited amount of thermal hysteresis data measured at different concentrations, mutants following the same methodology as our mutational analysis were utilized for data augmentation. A regression model was trained on these mutants with artificially attributed activity values. The values were determined based on associated activity seed protein activity value and the predictions provided from our ice binding model. These mutants were derived from anchor proteins, which are close to the seed proteins in ProtBERT space and screened to be above a 0.6 BLAST identity fraction threshold (**Figure 4E**). The 8 activity seed proteins derived from literature along with their mutation-targeted amino acid (alanine/threonine) are listed in **Supp. Table 1**. Note that only 6 activity seed proteins have anchor proteins due to the selection criteria. The model was then tested and evaluated with proteins that were experimentally tested for activity. As shown in **Figure 4F**, the initial model predictions displayed high performance (mean R^2^ score of 0.67), with 6 out of 8 seed proteins achieving an R^2^ score of 0.79 or above.

Besides assessing the model quantitatively with R^2^, we also assessed the model qualitatively by using our ice binding data corpus as an additional testing set. The objective is to assess the ability of the model to determine logical activity trends based on ice binding potential and concentration of the proteins. The results shown in the plot on **Figure 4F** indicates that the model has acquired a logical context for the activity trends. There is a significant difference in the activity value distributions between ice binding and non-ice binding proteins. The non-ice binding proteins maintain activity values near zero, which is the expected outcome. (Non-ice binding proteins should not have ice binding activity.) On the other hand, the activity values for ice binding proteins are significantly higher, and the values increase, as concentration increases, which fit our initial hypotheses. While the model may not display outstanding quantitative performance given by the R^2^ metrics, the model does provide adequate qualitative context to make adequate comparative predictions.

## Discussion

The development of a robust and scalable pipeline for predicting ice binding, expression, and activity from protein sequences represents a significant advancement in the field of protein engineering for cold environments. Our findings demonstrate that a harmonized and curated data corpus, when coupled with state-of-the-art protein language models, can effectively drive predictive tasks previously constrained by sparse, inconsistent annotations and limited training data.

Our initial contribution lies in assembling a large and functionally consistent dataset for ice binding proteins. Prior datasets were orders of magnitude smaller and often limited by ambiguous annotation or low diversity. By applying a controlled vocabulary across public databases and carefully filtering potential data leakage, we curated over 39,000 protein sequences, enabling us to train models that generalize beyond narrow functional motifs or taxa. The success of this harmonized corpus is evidenced by our sequence-based model’s performance, achieving accuracies over 89% on independent and held-out datasets, surpassing many traditional machine learning approaches in both scale and specificity.

While the accuracy scores were certainly comparable to those seen in previously published models, we wanted to test if our model contains the necessary biological context for determining the ice binding potential of a protein (3, 4, 5, 6). To this end, we engineered small but significant perturbations in its sequence inputs via targeted mutational analysis. By systematically mutating threonine and alanine residues—amino acids known to play critical roles in ice binding—we observed sensitivity patterns that aligned with current literature. The model’s ability to distinguish between threonine- and alanine-dependent binding motifs within different proteins suggests that it captured context-specific determinants of ice binding, not just global sequence features. Interestingly, the model’s predictions diverged from ESM2’s stability-based scoring in certain cases, underscoring a key distinction: while ESM2 provides insight into mutation favorability from an evolutionary perspective, our fine-tuned model directly encodes task-specific function. The divergence between these models also highlights the complexity of the ice-binding phenotype, where functional potential may not always correlate with evolutionary favorability or structural stability. These findings suggest that combining models trained on orthogonal objectives can provide complementary insights into protein design and function.

In addition to binding potential, our study tackled the important bottleneck of protein expression, a critical step in translating *in silico* predictions into experimentally tractable candidates. Using our engineered *Pichia pastoris* system, we trained an expression model that achieved decent performance (R^2^ = 0.64) during initial testing, with low false negative rates for highly expressed proteins. However, performance varied across sub-libraries. The INP sub-library, for instance, showed the highest correlation (0.75), while the AFP and P19614 sub-libraries exhibited lower predictive power. We attribute these discrepancies to training data limitations and the intrinsic diversity of the AFP family, which encompasses a broad range of structural and functional motifs. Our deeper subgroup analysis within the AFP library revealed specific seed groups that consistently underperformed, identifying potential targets for further refinement or data augmentation.

Lastly, our attempt to model ice activity from sequence, while nascent, opens promising avenues for engineering functional improvements beyond binary binding predictions. By leveraging model-driven mutational data for regression, we initiated a framework for associating sequence variation with thermal hysteresis—a more nuanced functional readout of antifreeze activity. Based on our results, the model performed adequately in some groups of proteins while struggling in others. This is due to data scarcity, as the groups of proteins the model struggled with had no associated training data context. However, a test screen of the ice binding dataset revealed significant differences between the predictions of the ice binding and non-ice binding proteins in a concentration-dependent manner, indicating that the model has developed an adequate knowledge base to potentially discern ice binding activity. Overall, the approach points toward the feasibility of integrating sequence space exploration with predictive functional scoring to prioritize high-performance candidates.

Several limitations merit discussion. While our models demonstrate strong generalization within known data, performance on entirely novel protein folds (*i*.*e*., chimeras) or environmental contexts remains untested. However, our model’s framework allows for tailoring to specific user-defined context (*i*.*e*., expression host). This should be seen as a warning prior to use, as the published model has not been trained on those novel protein folds. Additionally, while our mutational analyses provide interpretability, they remain computational and should be validated experimentally. For example, the training data for our activity model were derived from synthetically generated mutant proteins based on known ice binding proteins. Future work will incorporate high-throughput functional assays and directed evolution to close the loop between prediction and validation.

In conclusion, this study presents an integrated platform that moves beyond traditional discovery paradigms by combining scalable data curation, powerful sequence embeddings, mutational interpretability, and host-aware expression modeling. These tools collectively enable more targeted and rational engineering of ice binding proteins, which hold potential across cryopreservation, climate-resilient agriculture, and industrial biotechnology. As our dataset and model repertoire continue to expand, so too will our capacity to understand and manipulate the molecular basis of ice-protein interactions.

## Materials and Methods

### Ice Binding Metagenomic Library Curation (1st and 2nd Generations)

The 1st generation ice binding metagenomic library was designed using a jackHMMER search of public and proprietary databases. The input sequences for jackHMMER (*i*.*e*., metagenomic seeds) included 10 AFPs and 2 INPs (**Supp. Table 2**). (Note that metagenomic seeds are different from the activity seed proteins used in the activity model training explained below.) Metagenomic seeds were selected to optimize for structural and taxonomic diversity, and all seeds have biochemical evidence of ice binding activity in the literature. The jackHMMER search identified over 80,000 homologs. Of these homologs, fragments and sequences with invalid amino acids were filtered out, as well as sequences with a bitscore < 0.3 and sequences of length (0.5x < seed (x) < 1.5x). Approximately 1300 AFP and INP candidates were selected for the final 1st generation metagenomic library using an in-house multifactorial sampling which maximized the diversity of protein profiles with respect to protein size, similarity to known seeds, length and density of beta sheets in the secondary structure. Secondary structure composition was determined from predicted structures using DSSP (Dictionary of Secondary Structure of proteins), which uses hydrogen-bonding distances between main-chain atoms in a pdb file to assign residues to one of three secondary structure categories: helix, beta sheet, and loop (22). We then normalized by the length of the protein to determine the overall percent composition. To improve DNA synthesis success for the selected AFP and INP genes, which are often repetitive and difficult to produce, a software pipeline minimized self-homologies during encoding of the selected proteins into nucleotide sequences (23).

A 2nd generation metagenomic library was designed with the goal of optimizing the ice binding protein candidate pool. 2nd generation protein candidates were identified using a jackHMMER search of public and proprietary databases, where proprietary databases were expanded by 530 million protein sequences as compared to the 1st generation library, for a total of 3.5 billion sequences. The metagenomic seeds for the 2nd generation jackHMMER search included 15 AFPs and 2 INPs (**Supp. Table 2**), and these seeds were selected by re-using 1st generation metagenomic seeds and library hits that exhibited successful expression and ice binding activity. New seeds were also selected to increase structural diversity of the candidate pool compared to the 1st generation metagenomic library. In addition to the jackHMMER homology search, the 2nd generation metagenomic seeds were used to run a FoldSeek search of the AlphaFold database. The combined sequence and structure-based search identified over 160k homologs, from which 664 AFP and INP candidates were selected for the final 2nd generation metagenomic library. The final metagenomic library candidates were selected by first filtering out fragments, sequences with invalid amino acids, sequences with a bitscore < 0.3, sequences outside of length (0.5x < seed (x) < 1.5x), and percent identity thresholds to metagenomic seeds above 20%, and proteins not classified as BSL1. Additional candidates were filtered out using the ice binding model described here, where sequences classified as ‘non-ice binding’ were omitted. The final 2nd generation library candidates were diversified by selecting homologs for each of the 17 seeds. Homologs were then downsampled using multiple parallel methods: i) multifactorial sampling was used to maximize diversity based on protein size, sequence-similarity to known ice binding proteins, structural alignment to the metagenomic seed structure using root mean square deviations (RMSDs) calculated by US-align and secondary structure composition using the methods described above; ii) sequences were selected with the highest predicted value from the expression model reported here; iii) sequences were selected with the highest confidence for ice classification using the stability analysis reported here; iv) candidates were sampled with the lowest standard deviation for ice relevance (24).

### Measuring Protein Expression in *Pichia pastoris* with the HiBiT Assay

The construct design included an N-terminal secretion signal, ice binding protein of interest, TEV protease cleavage site, Strep-II affinity tag for purification, a flexible linker, and HiBiT tag for detection in the mentioned order. For the first generation library, 1033 AFP constructs and 274 INP constructs were synthesized through Twist Biosciences or internally at Ginkgo Bioworks for difficult templates. For the second generation library, a total of 1053 constructs were synthesized, including 492 AFPs and 111 INPs from metagenomic sourcing. These constructs were linearized and integrated as a single-copy into an internally identified locus in *P. pastoris*. 4 clones per construct were advanced for expression testing. Two base strains with different genetic backgrounds were tested and the better expressor was chosen for further strain build and test cycles.

Following strain engineering, strains were cultivated in 384-well plates for high-throughput expression testing. Briefly, cultures were grown in minimal media for induction of protein expression. After growth and induction, supernatants were collected and used as an input for the HiBiT detection assay which was executed using the manufacturer’s protocol (Promega Corporation). An in-plate positive control strain was used to normalize expression across different plates in the high-throughput screen. A negative control wild-type *P. pastoris* strain that does express any HiBiT-tagged protein was also present in each cultivation plate.

The QC evaluation of each plate in the assay was based on the coefficient of variation (CV) and Z’ of the HiBiT luminescence of the positive and negative control strains in each plate. CV is a measure of measurement repeatability, across all replicates of the positive control within each assay plate, based on the measured quantity value (HiBiT-tagged protein concentration). Assay containers with a CV value above 40% will not pass QC. The Z’ (Z-prime) is used to assess resolution between the positive and negative controls. A Z’ greater than 0 indicates that the assay has suitable resolution between the positive and negative controls for screening. In addition, in each plate a calibration curve is made using a known-concentration standard solution of purified recombinant 36kDa HaloTag® protein (Promega Corporation) to enable calculation of relative HiBiT-tagged protein concentrations in the measured samples. The R^2^ of linear regression fit of the standard curve is another QC metric obtained in the assay (R^2^ < 0.95 will not pass QC). >96% of the screened plates passed QC in the screens.

Using the HiBiT assay, the library was ranked for expression. HiBiT blotting was executed on a subset of samples to confirm protein size and integrity, also using the manufacturer’s protocol (Promega Corporation).

### Measuring Activity with Thermal Hysteresis Assay

Samples of AFPs were obtained from Ginkgo Bioworks and tested using a nanoliter osmometer described in literature (25, 26, 27). Briefly, a nanoliter amount of the sample was injected into an oil droplet held by a copper disc, which was placed on a temperature-controlled stage. The sample was then cooled to −30 °C to ensure complete freezing. The temperature of the sample was then slowly increased to melt most of the bulk ice and to obtain a single crystal of ~15 µm in diameter. This was achieved by careful temperature alterations and observation via a USB camera mounted on an E200 Nikon microscope (Nikon, Japan). After a single crystal at the desired size was obtained, the melting point of the crystal was documented, and the temperature was slightly decreased by 0.1 °C to prevent premature melting. At this point, a minute annealing time was initiated to allow the AFPs to bind to the surface of the ice crystal. After the annealing period was up, the temperature was decreased at a fixed rate of 0.15 °C/min until a growth burst was observed, which was defined as the non-equilibrium freezing point. The thermal hysteresis (TH) activity is defined as the difference between the melting point and the non-equilibrium freezing point of the crystal in °C.

### Expression and Activity Data Preprocessing

The expression model dataset comprises data from the 1st generation metagenomic library based on the HiBiT expression assay. In total, 833 datapoints were provided for training. Prior to training, the expression data was log normalized and stratified into low and high expression based on the threshold of 0.25, as shown in **Figure 2C**.

The initial protein activity dataset consisted of 8 datapoints (referred to as ‘activity seed proteins’) acquired from literature (**Figure 4E**). Protein activity refers to the thermal hysteresis activity that a protein is able to confer. This is determined from the thermal hysteresis assay from an experimental context. The associated activity seed protein concentration (in μM) during the activity measurement is also captured. Due to the limited nature of the dataset, data augmentation was implemented utilizing the mutation generation strategy explained below. The process of data augmentation is illustrated in **Figure 4E**. First, a set of 52 ‘anchor proteins’ were curated to serve as the basis of the generation of the mutants. The 52 anchor proteins were determined through ProtBERT analysis, where these proteins were selected based on distance to the seed proteins in ProtBERT embedding space. These anchor proteins were then screened through BLAST with an identity fraction threshold of 0.6. (Note that only 6 activity seed proteins have anchor proteins based on this criteria.) The anchor proteins were mutated based on the amino acid sensitivity of their associated seed proteins (determined from our mutational analysis). The 6 activity seed proteins and their mutation-targeted amino acids are listed in **Supp. Table 1**. After generating mutants of the anchor proteins and attributing appropriate activity values as so: (activity seed protein activity value) * (ice binding model prediction of mutant), these mutants were then aggregated into a training set (following a 30-70 test-train split). The associated concentration from the seed proteins is also attributed to the mutants. In total, there are approximately 200,000 mutants and 900,000 unique activity-concentration datapoints in the initial training set. For additional verification, subsets of this training set were created based on the mutation percentage threshold, depicted in **Figure 2D**. (For example, a subset with a 20% mutation percentage threshold contains all mutants with at least 20% of their alanines or threonines mutated.) The thresholds were set from 10 to 90% in 10% increments. BLAST alignment analysis was also conducted for each subset by aligning mutant sequences to their associated activity seed protein sequences. The average identity fraction for each subset was calculated and used as an objective measure for the sequence similarity between the training subsets and the testing set (activity seed proteins). These subsets were used to train additional models to compare performance metrics with the initial model, shown in **Supp. Figure D**.

### Model Perturbation Processes

In order to perform a stability analysis on the ice binding model, Gaussian noise was introduced prior to the fine-tuning layers of the model, as shown in **Figure 3A**. For the injection of Gaussian noise, a simple Gaussian filter with a mean of 0 and a sigma varying from 0 to 0.2 was applied to add a random amount of Gaussian noise to all of the ProtBERT embeddings. (The embeddings were already normalized to a scale of −1 to 1.) Resultant changes in model probabilities were then calculated and computed as the standard deviation of probabilities for each protein. This was conducted for 10 iterations to produce the average standard deviation. Note that lower average standard variations indicate higher prediction stability (less variation in model probabilities). Results are shown in **Supp. Figure B**.

In order to introduce biological noise to the model, the mutational generation workflow was applied. Based on literature guidance, certain amino acids were determined to be critical to ice binding function (*i*.*e*., alanine, threonine). Mutants were then generated from these ice binding proteins by mutating those critical amino acids into another amino acid at random (**Figure 3C**). To determine the sensitivity of the model to the number of mutations, mutants with differing numbers of mutations from 1 to the maximum number of mutations possible for each protein were produced. (The number of mutations refers to the number of a certain critical amino acid that were mutated per mutant, as shown in **Figure 3C**.) The number of mutations was then normalized by the total number of alanines in the original protein for the alanine mutated number percentage for each mutant. (For example, mutating all of the alanines present in the original protein would yield a mutated number percentage of 100%.)

### Model Architectures and ProtBERT

All of the models introduced in this manuscript utilize ProteinBERT (ProtBERT) as the base model (**Figure 1**). ProtBERT is a deep learning masked language model that takes protein amino acid sequences and outputs a set of 1024-dimensional embeddings that is used for downstream protein predictors. As shown in **Figure 1**, these embeddings were then fed to separately trained model heads to predict the ice binding potential (the likelihood of a protein to have ice binding function), ice function based on GenBank data (the likelihood to have an antifreeze, ice-binding, ice nucleation, or ice structuring function annotation), ice expression (predicting the protein expression level based on the HiBiT assay), and ice activity (predicting the protein activity level based on the thermal hysteresis assay).

**Figure 1:**
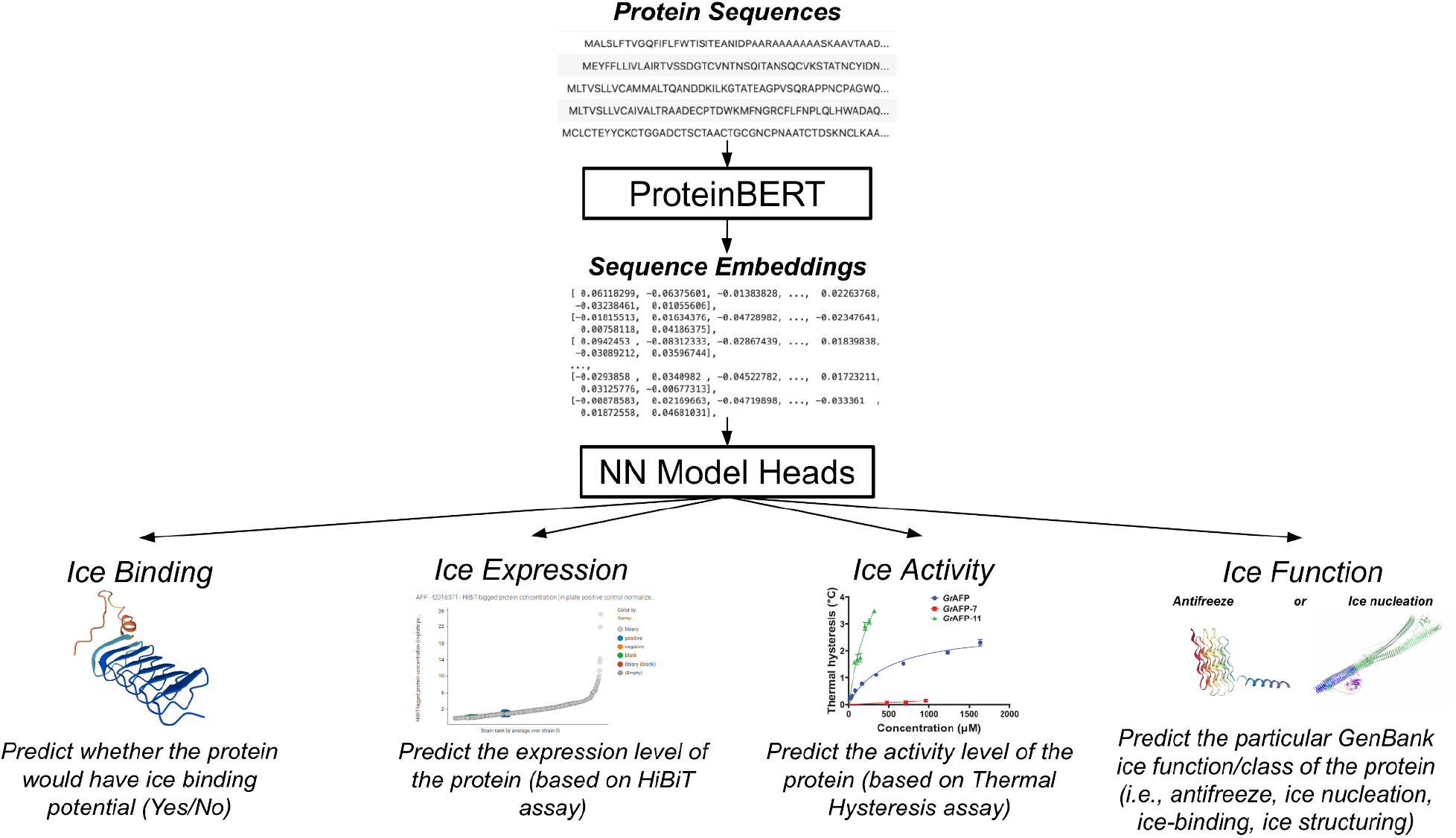
General overview of the ICE models for predicting ice binding, function, expression, and activity. The models all use ProtBERT as the base, which intakes protein sequences to output corresponding vector embeddings. These embeddings will then be fed to neural network model heads for predicting various aspects of ice proteins, including ice binding potential (whether a protein can be classified as an ice binding protein), ice expression (determining the HiBiT expression level of the protein within a Pichia host), and ice activity (determining the protein activity based on the thermal hysteresis assay). A model has also been developed to predict ice function (whether a protein is classified as having a particular GenBank ice function: antifreeze, ice-binding, ice structuring, ice nucleation), which is elaborated in the Supplementary Information.

**Figure 2:**
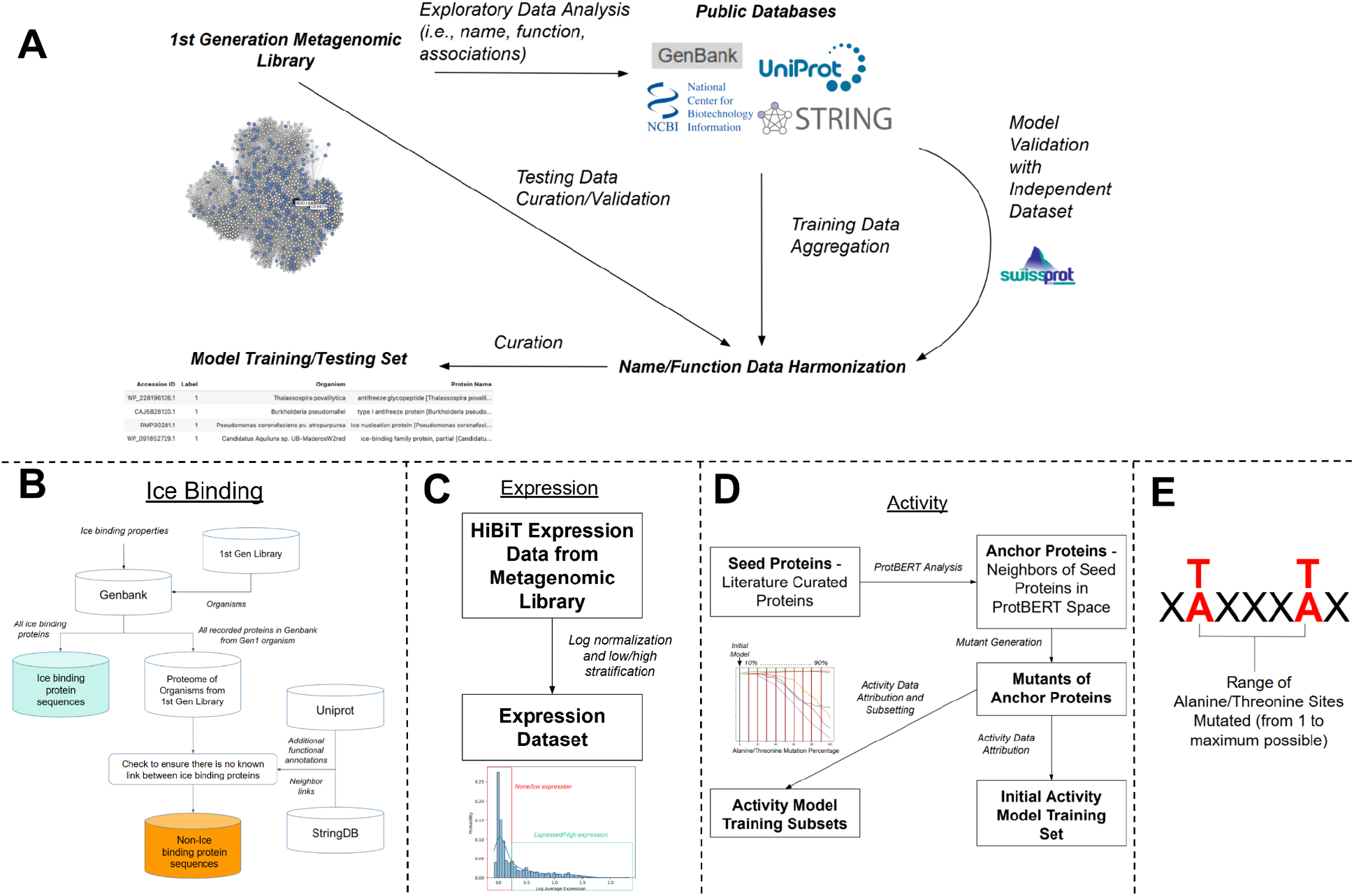
Training Data. A) General data curation process. The 1st generation metagenomic library is first screened through public databases for common protein names and functions. These annotations were then used to acquire all proteins from GenBank that share the same annotations. This expanded set of proteins was curated by obtaining their sequences, names, source organisms, and other available data from the databases. This ultimately becomes the training dataset for the ice binding model. Additional testing/validation datasets were curated from the original metagenomic library and independent datasets. The data is normalized and harmonized for consistency prior to model training and testing. B) Data curation for the ice model heads. The ice binding model utilizes both ice binding and non-ice binding protein sequences. The curation of ice binding sequences is elaborated in A. For sourcing non-ice binding proteins, the proteomes of the organisms sourcing the 1st generation metagenomic library were obtained, and a number of random proteins equal to the number of ice binding proteins were chosen. These non-ice binding proteins were verified to have no known links to the ice binding proteins, assuring that there is no leakage between the groups. C) Training corpus curation for the ice expression model. The training corpus for the expression model head was curated from the HiBiT expression data from the metagenomic library. The data was then log normalized and stratified based on the data distribution. D) Training set curation for the Ice activity model. The training set for the activity model head was curated from a set of seed proteins (proteins with literature activity data). A set of anchor proteins were then derived from the list of seed proteins if they were neighbors to the seed proteins in ProtBERT space (screened based on the distance determined from the vector embeddings). Mutants were generated from the seed and anchor proteins by mutating their alanines or threonines to a random amino acid (as shown in E). These mutants would then be used for the initial training set. Further analysis was conducted with the activity model by subsetting the initial training set based on the mutation percentage threshold, depicted in the representative plot. (For example, a subset with a 20% mutation percentage threshold contains all mutants with at least 20% of their alanines or threonines mutated.) E) Mutation generation schematic. For the mutational analysis, ice binding proteins were mutated at their alanine or threonine sites. The mutations involve mutating the alanine/threonine to a random amino acid (termed alanine/threonine-to-random mutations). Various mutants were created containing a varying number of alanine/threonine-to-random mutations from just 1 alanine/threonine mutated to the maximum number of alanines/threonines mutated.

For the ice binding model head (**Figure 3A**), a dense neural network was trained to take the 1024-dimensional embeddings to output a label signifying ice binding or non-ice binding. There are 5 hidden layers with 512, 256, 128, 64, and 32 neurons, respectively. Each layer is batch normalized, and a rectified linear unit (ReLU) activation function is applied to the final hidden layer. A sigmoid activation function is used in the final output layer to output a probability value from 0.0 to 1.0. A cutoff of 0.5 was instituted for the binary labels ice binding and non-ice binding.

For the ice expression model head (**Figure 4A**), a dense neural regressor network was trained to take the embeddings to output a final expression value that would have been obtained through the HiBiT assay in a *Pichia* host. There are 5 hidden layers with 512, 256, 128, 64, and 32 neurons, respectively. Each layer is batch normalized, and a rectified linear unit (ReLU) activation function is applied to the final layer, which outputs the predicted expression value. Because the initial expression data was natural log normalized, the output of the model will be a log normalized value.

**Figure 3:**
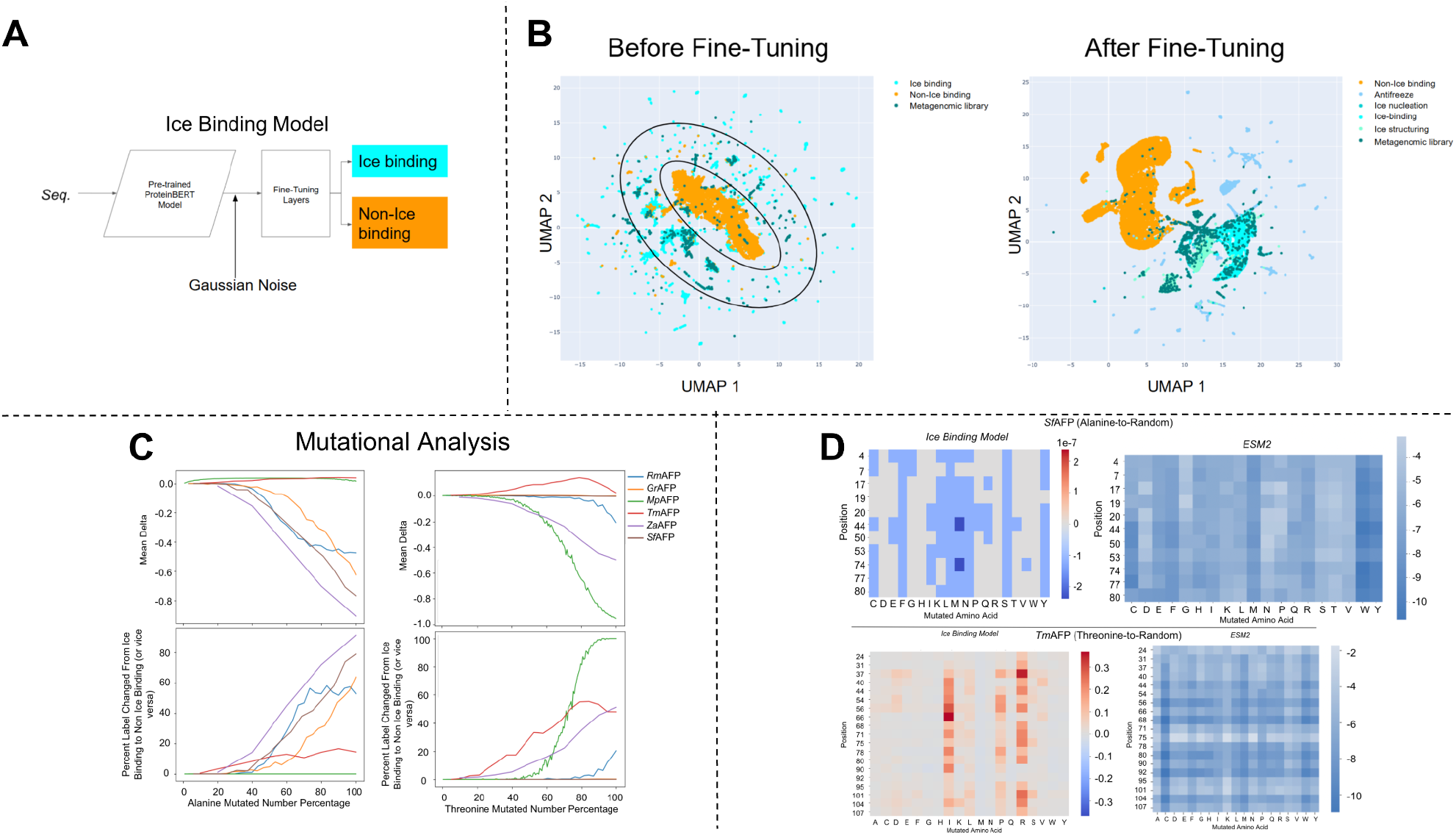
Ice Binding Model and Associated Analysis. A) Architecture of the ice binding model. For stability analysis, Gaussian noise is inserted between ProtBERT and fine-tuning layers. B) UMAP plots of the high dimensional embedding space before and after the ice binding model fine-tuning. Before fine-tuning, ProtBERT already partially segregated the proteins by ice binding potential, forming a dominant cluster of non-ice binding proteins (in orange) in the center and a ring of ice binding proteins (in blue and dark green) around that central cluster. After fine tuning with the ice binding modeling layers, the clustering of non-ice and ice binding proteins is more distinct. (Left cluster = non-ice binding; right cluster = ice binding) Ice binding proteins were further classified into their ice functions as illustrated in the UMAP. C) For mutational analysis, the model predictions of the produced synthetic mutants were compared to that of their original proteins (shown under ‘Accession ID’) to calculate the mean probability deltas. The mutants have a varying number of their alanines or threonines mutated, as depicted in the mutated number percentage. As shown in the plots, mutants generally exhibit a decrease in their prediction probability, indicating that the mutations are influencing the ice binding model’s predictions and increasing the number of changed prediction labels (going from predicting ice binding to non-ice binding). Moreover, different proteins respond to the different amino acid mutations (alanine/threonine) differently, as shown by comparing results from alanine mutations (left column) and threonine mutations (right column). D) Representative comparison of results between the ice binding model presented in this paper and the ESM2 model for single alanine/threonine-to-random mutations. The results from the ice binding model correspond to the probability deltas due to the mutations. The results from ESM2 correspond to their protein stability metric. Blue indicates a negative probability delta or protein stability metric, while red indicates an increase. While the two models generally show similar trends in their respective metrics for alanine-to-random mutations, they tend to differ for threonine-to-random mutations, as indicated by the increase in ice binding potential for the mutants as opposed to the decrease in protein stability metric.

For the ice activity model head (**Figure 4D**), a dense neural network model was developed to predict the thermal hysteresis activity of proteins for any given concentration in μM. The model takes in the associated embeddings corresponding to the protein first. The first part of the neural network contains 4 hidden layers with 512, 256, 128, and 64 neurons, respectively. It outputs a 32-dimensional embedding which is then aggregated with the base-10 log of the input protein concentration. The aggregate is then fed into a regression node containing 1 hidden layer of 16 neurons. A ReLU function is applied to every hidden layer in the network. The final output contains the predicted activity value of the protein at the provided concentration.

**Figure 4:**
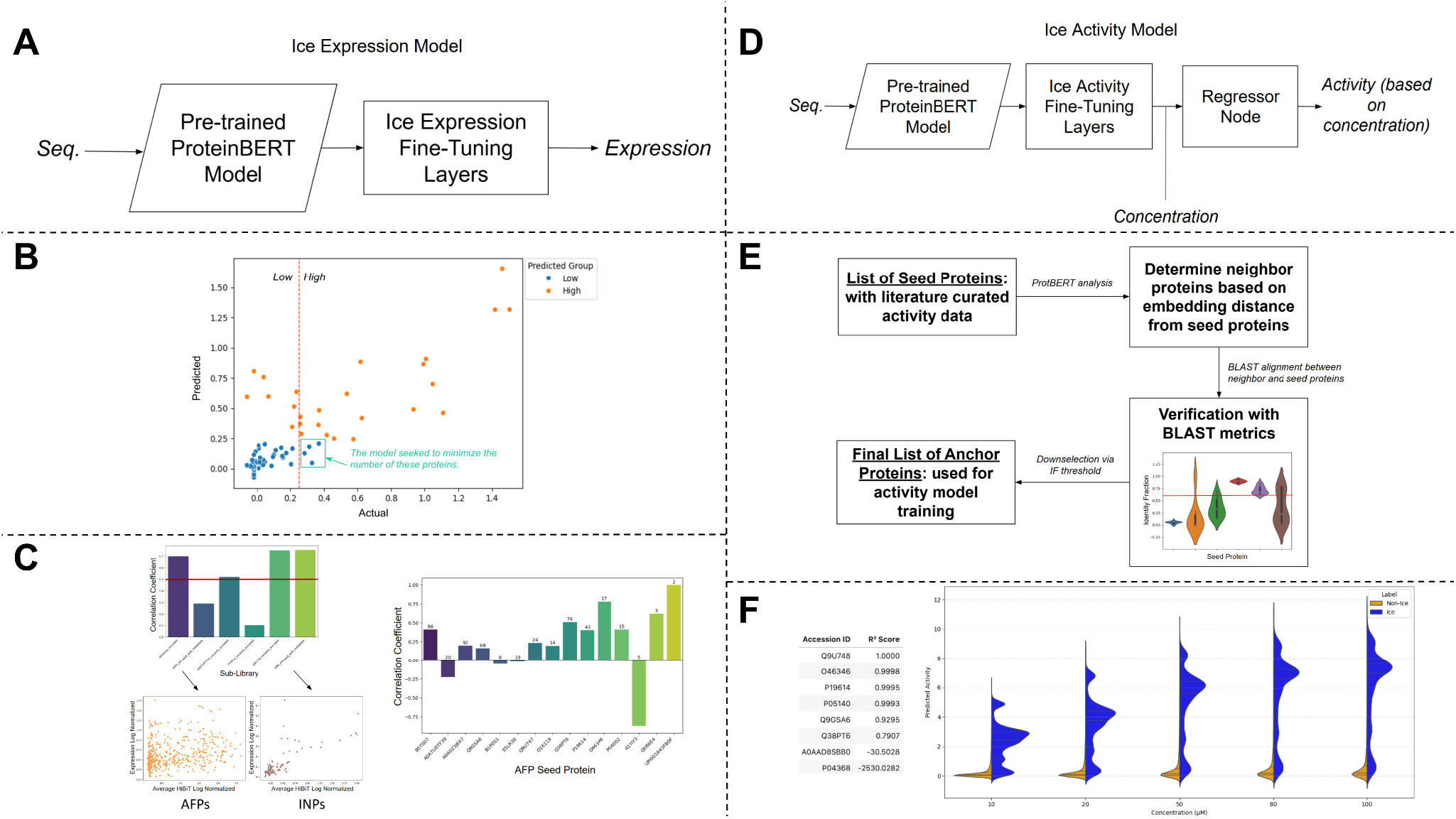
Results from the Ice Expression and Ice Activity Models. A) Architecture of the ice protein expression model. The final ice expression model utilizes a simple regression head for its fine-tuning layers to output a predicted HiBiT expression value from the *Pichia* host. B) Expression model results. The scatter plot depicts the low vs high expressed proteins and the predicted group by the model (blue vs orange). The green box depicts the number of proteins considered as false negatives, and the model was optimized to minimize that number. The simple regression head also minimizes the false negative, which is important when selecting for highly expressed proteins. C) 2nd generation metagenomic analysis for the expression model. Left: After conducting a blinded validation of the 2nd generation metagenomic library on the expression model trained on the 1st generation metagenomic library, the correlation between the predicted expression values and the experimental values for each sub-library. The representative scatter plots below depict the low and high correlation seen from the aggregate results above. The model achieved a correlation of over 0.5 for 4 of the 6 sub-libraries. Right: The correlations for the AFPs grouped by their sourced seed protein were determined and shown in the plot. The number above each bar indicates the number of AFPs in each group. D) Architecture of the ice protein activity model. A simple regression model was trained to predict thermal hysteresis activity of proteins. The ProtBERT embeddings were first passed into the fine-tuning layers. The concentration of the protein (in μM) was then input before a regressor node for the prediction. Therefore, the predicted activity is dependent on the concentration of the protein. E) Activity Model Anchor Protein Curation Methodology. Using the list of seed proteins, or proteins with available activity data determined from literature, ProtBERT analysis was conducted to determine the neighbor proteins. These neighbor proteins are close to the seed proteins in ProtBERT embedding space, which is dependent on protein sequence. For further verification, these neighbor proteins were then passed through BLAST for alignments with their associated seed proteins. The plot shows a representative set of neighbor proteins and their identity fraction in relation to their associated seed protein. The red line denotes the 0.6 identity fraction threshold. This threshold is used to down-select the neighbor proteins to ensure similarity in sequence space for downstream activity data attribution. The final list of anchor proteins will be used for synthetic data augmentation to form the training corpus for the activity ranking model. F) Performance metrics of the activity model. The data table on the left displays the R^2^ scores for each seed protein after initial activity model testing. The plot on the right compares the activity model predictions for the ice binding data corpus, separated by ice binding and non-ice binding proteins, across varied concentrations. The plot shows that the model outputs significantly different predictions for ice binding and non-ice binding proteins, indicating that it can distinguish between the two groups of proteins. The non-ice binding proteins receive predictions near zero activity, while the ice binding proteins display increasing activity values as concentration increases.

### Model Training and Accuracy Testing

For training of all model heads, the weights of the pre-trained ProtBERT model were frozen. Each model head then undergoes training based on specific sets of training parameters and hyperparameters. All models used the Adam optimizer to minimize the loss function with a learning rate of 0.001. Datasets are split for 70% training and 30% testing except for the activity model, which utilized an augmented dataset for training and literature-tested protein data for testing. Early stopping was employed for training models with a patience of 15 epochs to prevent overfitting.

For both the ice binding and ice function models, the binary cross entropy loss function was used to optimize the model weights, a batch size of 512 datapoints was used for training, and accuracy was determined using the percentage of correctly predicted labels. For the ice expression model, the MSE loss function was used with a batch size of 64. Accuracy was determined by the R^2^ between the predicted expression value and actual value. Prior to developing the model, the HiBiT expression data was first log normalized to reduce the skewness of the data distribution. The data was then segregated into two groups, with expression below 0.25 is considered ‘low to no expression’, while the expression above 0.25 is considered ‘high expression’. The data was stratified based on these expression groups to balance the training dataset. For the ice activity model, the MSE loss function was used, and a batch size of 2048 was employed. Protein concentrations were log normalized prior to training. Accuracy was determined by the R^2^ score between the predicted activity value and the actual literature value at the same protein concentrations. For testing, the seed protein activity values measured at different concentrations were used. The ensemble prediction (taken from the best model trained at each cross-validation fold) was used during testing. Training subsets were also formed to assess potential data leakage. A qualitative assessment was also conducted by using the ice binding data corpus as a testing set to determine the trends between the predicted activity values for the ice binding vs non-ice binding proteins from the corpus. Concentrations of 10, 20, 50, 80, and 100 μM were arbitrarily chosen to mimic potential tested protein concentrations.

## Data and Toolkit Sharing Plans

All data used to train the models along with a toolkit armed with the models can be found in this repository: https://github.com/netrias/ICEPIC. The repository includes a set of instructions on how one can use the toolkit to predict whether a new protein sequence is ice binding, what its expression level in the P. pastoris host, and its activity level as assessed by thermal hysteresis. If you are working with a different host or activity measurement, please email the corresponding authors to obtain guidance on fine-tuning the toolkit for your needs.

## Acknowledgements

This work was supported by the Defense Advanced Research Projects Agency (DARPA) under contract number 11844 as part of the project titled *Integrated Computational Exploration of Proteins for Ice Control*. The views, opinions, and findings expressed are those of the authors and should not be interpreted as representing the official views or policies of the U.S. Government. The authors would also like to thank Dr. Ran Drori from Yeshiva University for his expertise and guidance on producing this manuscript.

## References

1. Bar Dolev M, Braslavsky I, Davies PL. Ice-Binding Proteins and Their Function. Annual Review of Biochemistry. 2016;85(Volume 85, 2016):515–42.

2. Box ICH, van der Burg KRL, Marshall KE. Analysis of Ice-Binding Protein Evolution. In: Drori R, Stevens C, editors. Ice Binding Proteins: Methods and Protocols. New York, NY: Springer US; 2024. p. 219–29.

3. Dhibar S, Jana B. Accurate Prediction of Antifreeze Protein from Sequences through Natural Language Text Processing and Interpretable Machine Learning Approaches. The Journal of Physical Chemistry Letters. 2023;14(48):10727–35.

4. Usman M, Khan S, Lee J-A. AFP-LSE: Antifreeze Proteins Prediction Using Latent Space Encoding of Composition of k-Spaced Amino Acid Pairs. Scientific Reports. 2020;10(1):7197.

5. Ali F, Akbar S, Ghulam A, Maher ZA, Unar A, Talpur DB. AFP-CMBPred: Computational identification of antifreeze proteins by extending consensus sequences into multi-blocks evolutionary information. Computers in Biology and Medicine. 2021;139:105006.

6. Qi D, Liu T. VotePLMs-AFP: Identification of antifreeze proteins using transformer-embedding features and ensemble learning. Biochimica et Biophysica Acta (BBA) - General Subjects. 2024;1868(12):130721.

7. Kozuch DJ, Stillinger FH, Debenedetti PG. Combined molecular dynamics and neural network method for predicting protein antifreeze activity. Proceedings of the National Academy of Sciences. 2018;115(52):13252–7.

8. Verbitsky A, Boutet P, Eslami M. Metadata Harmonization from Biological Datasets with Language Models. bioRxiv. 2025:2025.01.15.633281.

9. Kandaswamy KK, Chou K-C, Martinetz T, Mller S, Suganthan PN, Sridharan S, et al. AFP-Pred: A random forest approach for predicting antifreeze proteins from sequence-derived properties. Journal of Theoretical Biology. 2011;270(1):56–62.

10. Zhao X, Ma Z, Yin M. Using support vector machine and evolutionary profiles to predict antifreeze protein sequences. Int J Mol Sci. 2012;13(2):2196–207.

11. Brandes N, Ofer D, Peleg Y, Rappoport N, Linial M. ProteinBERT: a universal deep-learning model of protein sequence and function. Bioinformatics. 2022;38(8):2102–10.

12. Garnham CP, Campbell RL, Davies PL. Anchored clathrate waters bind antifreeze proteins to ice. Proceedings of the National Academy of Sciences. 2011;108(18):7363–7.

13. Sun T, Lin F-H, Campbell RL, Allingham JS, Davies PL. An Antifreeze Protein Folds with an Interior Network of More Than 400 Semi-Clathrate Waters. Science. 2014;343(6172):795–8.

14. Ye Q, Eves R, Campbell RL, Davies PL. Crystal structure of an insect antifreeze protein reveals ordered waters on the ice-binding surface. Biochemical Journal. 2020;477(17):3271–86.

15. Liou Y-C, Daley ME, Graham LA, Kay CM, Walker VK, Sykes BD, et al. Folding and Structural Characterization of Highly Disulfide-Bonded Beetle Antifreeze Protein Produced in Bacteria. Protein Expression and Purification. 2000;19(1):148–57.

16. Scholl CL, Tsuda S, Graham LA, Davies PL. Crystal waters on the nine polyproline type II helical bundle springtail antifreeze protein from Granisotoma rainieri match the ice lattice. The FEBS Journal. 2021;288(14):4332–47.

17. Lin Z, Akin H, Rao R, Hie B, Zhu Z, Lu W, et al. Evolutionary-scale prediction of atomic-level protein structure with a language model. Science. 2023;379(6637):1123–30.

18. Liu M, Liang Y, Zhang H, Wu G, Wang L, Qian H, et al. Production of a recombinant carrot antifreeze protein by Pichia pastoris GS115 and its cryoprotective effects on frozen dough properties and bread quality. LWT. 2018;96:543–50.

19. Li Z, Xiong F, Lin Q, d’Anjou M, Daugulis AJ, Yang DSC, et al. Low-Temperature Increases the Yield of Biologically Active Herring Antifreeze Protein in Pichia pastoris. Protein Expression and Purification. 2001;21(3):438–45.

20. Kim EJ, Lee JH, Lee SG, Han SJ. Improving thermal hysteresis activity of antifreeze protein from recombinant Pichia pastoris by removal of N-glycosylation. Prep Biochem Biotechnol. 2017;47(3):299–304.

21. Loewen MC, Liu X, Davies PL, Daugulis AJ. Biosynthetic production of type II fish antifreeze protein: fermentation by Pichia pastoris. Applied Microbiology and Biotechnology. 1997;48(4):480–6.

22. Kabsch W, Sander C. Dictionary of protein secondary structure: Pattern recognition of hydrogen-bonded and geometrical features. Biopolymers. 1983;22(12):2577–637.

23. Zulkower V, Rosser S. DNA Chisel, a versatile sequence optimizer. Bioinformatics. 2020;36(16):4508–9.

24. Zhang C, Shine M, Pyle AM, Zhang Y. US-align: universal structure alignments of proteins, nucleic acids, and macromolecular complexes. Nature Methods. 2022;19(9):1109–15.

25. Braslavsky I, Drori R. LabVIEW-operated novel nanoliter osmometer for ice binding protein investigations. J Vis Exp. 2013(72):e4189.

26. Pariente N, Bar Dolev M, Braslavsky I. The Nanoliter Osmometer: Thermal Hysteresis Measurement. In: Drori R, Stevens C, editors. Ice Binding Proteins: Methods and Protocols. New York, NY: Springer US; 2024. p. 75–91.

27. Berger T, Meister K, DeVries AL, Eves R, Davies PL, Drori R. Synergy between Antifreeze Proteins Is Driven by Complementary Ice-Binding. Journal of the American Chemical Society. 2019;141(48):19144–50.

