## Supplementary Information for "ICEPIC: A Toolkit to Discover Ice Binding Proteins from Sequence"

Classification: Biological Sciences, Computational Biology (minor)

Keywords: Ice binding proteins, antifreeze proteins, ice nucleation proteins, protein engineering

** Co-corresponding Authors*
Mohammed Eslami,
Susan Buckhout-White,

**Supp. Figure A** displays the initial GenBank results of our exploratory data analysis of the 1st generation metagenomic library and development of a GenBank ice function model. **Supp. Figure AA** depicts plots of the names (left) and annotated functions (right) obtained from GenBank for the 1st generation metagenomic library. All names and functions present in the plots have appeared at least twice during the exploratory analysis. The significant points to note include the abundance of ice binding relevant terms in protein names. Approximately 70% of annotated names have ice functional annotations (*i.e.,* ‘ice-binding, ice nucleation, antifreeze, ice structuring). Note that the ‘ice-binding’ annotation in GenBank may be considered a subset annotation of the general category of ice binding proteins investigated in this manuscript. This became the basis for curating the proteins in our ice binding training corpus. Similarly, 41% of annotated protein functions are considered ice binding, and 40% of the functions are associated with the extracellular space, which reflects the nature of ice binding proteins. (They are located in the extracellular space.) A GenBank ice function model was developed (**Supp. Figure AB**) with a dense neural network was developed to predict whether a protein belongs in one of five classes derived from GenBank: antifreeze, ice-binding, ice nucleation, ice structuring, and non-ice. This model head consists of 7 hidden layers with 512, 256, 128, 64, 32, 16, and 8 neurons, respectively. Each layer is batch normalized, and a ReLU activation function was applied to the final hidden layer. A softmax activation function is applied to the final layer, which outputs the probabilities that input sequence belongs in each of the five aforementioned classes. The assigned function label of the input sequence is determined by the class with the highest probability. Testing the classification model for ice function yielded an accuracy of 90%. The confusion matrix of the true vs predicted classes is shown in **Supp. Figure AC** to identify specific classes that the model excels and struggles at predicting. Looking at true ice nucleation proteins, one of the classes that was delineated in the 1st generation metagenomic library, the model has a high accuracy of 98%. The model has a lower accuracy of 95% when capturing true antifreeze proteins. The high sequence and structural diversity of antifreeze proteins may be a cause for lower accuracy in predicting the antifreeze proteins.

**Supp. Figure B** displays the results of the stability analysis after Gaussian noise injection. Overall, the results indicate that the model is stable after Gaussian noise injection, with all of the proteins obtaining average standard deviations below the 0.5 threshold (used in the prediction of ice binding vs non-ice binding proteins). Moreover, the segregation of proteins based on their predictions reveals that proteins that were predicted as ice binding are generally more stable (lower average standard deviation) than those predicted as non-ice binding. This is indicative of the fact that the model is more certain of its ice binding predictions compared to its non-ice binding predictions. (This analysis assumed that all metagenomic library proteins are true ice binding proteins.) This is further supported from the UMAP plot on the left side of **Supp. Figure B**. This UMAP figure was taken from embeddings obtained in the fine-tuning layers of the ice binding model (**Figure 3B**). The figure illustrates the ice binding cluster on the right of the red-dotted line, while the non-ice binding cluster is illustrated to the left of the line. Proteins that are more stable are mostly located right of the line in the ice binding cluster. However, proteins generally become less stable as they get closer to the line boundary and into the non-ice binding cluster on the left. This corroborates with the conclusion that the model is highly certain of its ice binding (correct) predictions and less certain of its non-ice binding (incorrect) predictions, as shown in the distribution plot on the right side of **Supp. Figure B**.

**Supp. Figure C** includes supplementary information describing the activity model training dataset and subsets. The left plot depicts the distributions of the identity fraction of the anchor proteins grouped by activity seed proteins. These values were determined via BLAST pairwise alignments between each anchor protein and its associated activity seed protein in ProtBERT space. The distributions are bounded between 0.6 and 1 due to the threshold criteria implemented when selecting anchor proteins. The activity models were trained on mutants of these anchor proteins. Note that 6 of the original 8 seed proteins have anchor proteins due to the selection criteria. The right plot displays the resultant average identity fractions for each subset based on pairwise alignments between the mutants in the subset to their respective seed proteins. Generally, an increase in the percentage threshold used in forming the subset led to a decrease in the identity fraction. More mutations leads to more differences in sequence space, lowering the identity fraction. However, this is not necessarily a direct correspondence, as shown in the case for *P19614*, but an explanation is that the mutants were created from anchor proteins, not activity seed proteins. This was done to reduce data leakage between the training dataset (anchor mutants) and the testing dataset (activity seed proteins).

A potential concern to the high performance for certain seed proteins was the potential data leakage from the anchor protein mutants. Due to the mutant generation process, some mutants have few mutations, meaning that their sequences may be highly similar to those of the seed proteins. (ProtBERT is a sequence-based model, so the chosen anchor proteins would also have high sequence similarity (**Supp. Figure C**).) Therefore, the activity training dataset was subset based on the mutation percentage of the mutants. Mutation percentage thresholds of 10-90% in 10% increments were used to subset the data. Subsets also underwent BLAST alignment analysis comparing the mutants and their associated activity seed protein, and the average identity fraction for each subset was calculated. Models were then trained on the subsets with the same architecture and methodology, and the $R^2$ score for each protein is plotted against the average identity fraction of each trained subset (**Supp. Figure D**). As expected, the $R^2$ scores generally increased as the identity fraction increased, meaning that the models performed better the closer the training sequences were to the testing sequences (seed proteins). However, it should be noted that most high $R^2$ scores (near 1) were achieved with subsets having identity fractions around 0.7 to 0.85, meaning that despite the high model performance, the mutants were still substantially different to not induce significant data leakage. (For reference, an identity fraction of 0.9 indicates that at least 7 amino acids in the activity seed protein would have been mutated. The average number of amino acids needed across all activity seed proteins is 11.5.) The results also suggest that an identity fraction objective window (0.7 to 0.85) exists to maximize confidence in the model. Some discrepancies in the trends such as different performances for different activity seed proteins and presence of low $R^2$ scores for subsets with high identity fractions can be explained with **Supp. Figure C**. Proteins with lower identity fraction distributions (*i.e., Q38PT6*) had significant $R^2$ score decreases at lower percentage thresholds than proteins with higher distributions (*i.e., P19614*). The two proteins that displayed negative $R^2$ scores (*P04368, A0AAD8SBB0*) in the initial activity model had no anchor proteins (zero identity fraction), so the trained models had no context for these proteins. While the identity fraction was used as an objective measure to determine the effectiveness of subsetting the dataset in analyzing data leakage, it does not necessarily directly correspond to the mutation percentage threshold used for subsetting. (The mutants were created from anchor proteins, which are similar but not identical to the activity seed proteins in sequence space.) Overall, the activity model dataset did not significantly leak data when evaluating the test set.

**Supp. Figure E** depicts the results for our CD-Hit analysis on the ice binding and metagenomic library proteins in the ice binding model training corpus. CD-Hit is a tool used to assess sequence similarity by grouping similar sequences into clusters. The analysis was conducted at a 90% identity threshold. The analysis revealed 7959 different clusters from the ice binding and metagenomic library proteins (left plot). While this demonstrates that the corpus contains significantly similar sequences, it is still significantly higher than the size of the dataset used by other models in literature. A deeper analysis into cluster sizes revealed that a great majority (85%) of clusters contain only 1 or 2 sequences, illustrating that most of the corpus is unique. An analysis into the predominant labels for the clusters showed the different functions associated with each cluster (right plot). Notably, the number of clusters (2328) with the 'antifreeze' label far exceeds the 400-500 antifreeze protein sequences used by other models.

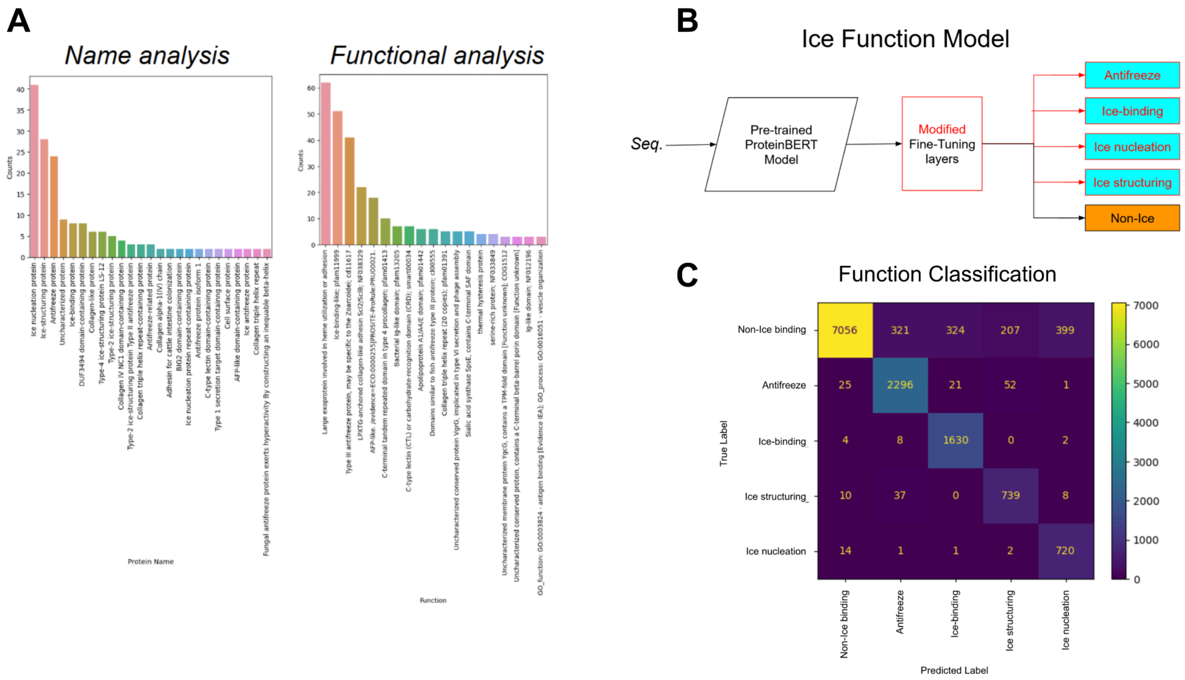


**Figure A. Plots of Protein Names and Functions from the 1st Generation Metagenomic Library Used to Develop the Ice Function Model.** A) The protein names and functions were obtained from GenBank for the metagenomic library proteins that have data in GenBank. The plots only includes names and functions that appeared more than once. B) Architecture of the ice function model. The ice function model architecture largely follows that of the ice binding model architecture. The red font in the architecture indicates the differences between it and the ice binding model architecture. C) The confusion matrix after classifying a held out dataset with original metagenomic library proteins, displaying the strong performance when classifying between the functional labels for ice binding proteins.

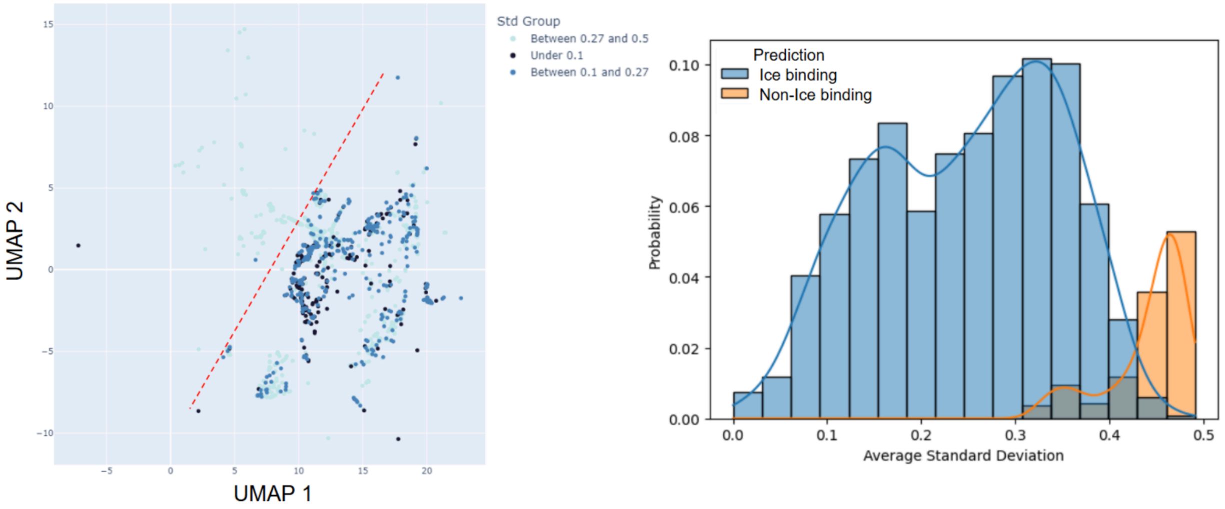


**Figure B. Stability analysis of the ice binding model via injecting Gaussian noise into the model.** The degree of stability is determined by the average standard deviation of predictions caused by the noise. The UMAP plot on the left depicts that the ice binding proteins in the ice binding cluster (right cluster) exhibit more prediction stability (low standard deviation) on average than the proteins found in the non-ice binding cluster (left cluster with higher standard deviation). The red dotted line shows the boundary between the two clusters. The distribution plot of the average standard deviation is shown on the right, further supporting the fact that false negatives (ice binding proteins predicted as non-ice binding) have higher instability. In other words, the model is more certain of the true positives and less certain about the false negatives.

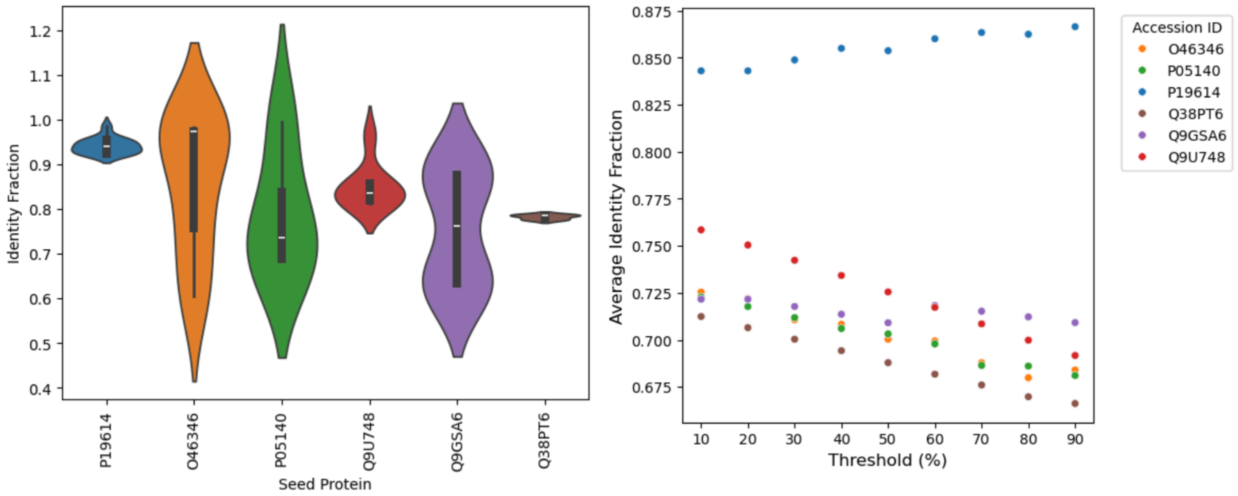


**Figure C. Supplemental analysis of the activity model training dataset and subsets.** Left: Identity fraction of anchor proteins compared to seed proteins via pairwise BLAST alignments between each anchor protein and its associated seed proteins. The plot depicts the distributions of identity fraction of the anchor proteins grouped by seed proteins. Right: Change in identity fractions of subsets based on mutation percentage threshold grouped by seed proteins. The subset thresholds were set by the mutation percentages of alanines/threonines, and each subset contains all mutants that meets the threshold in terms of mutation percentage. As indicated in the plot, the mutants generally experience lower identity fractions as the subset threshold increases, since more mutations lead to more sequence differences. However, it is noted that the mutants were derived from anchor proteins, not seed proteins, so mutations do not necessarily correspond to a decrease in identity fraction.

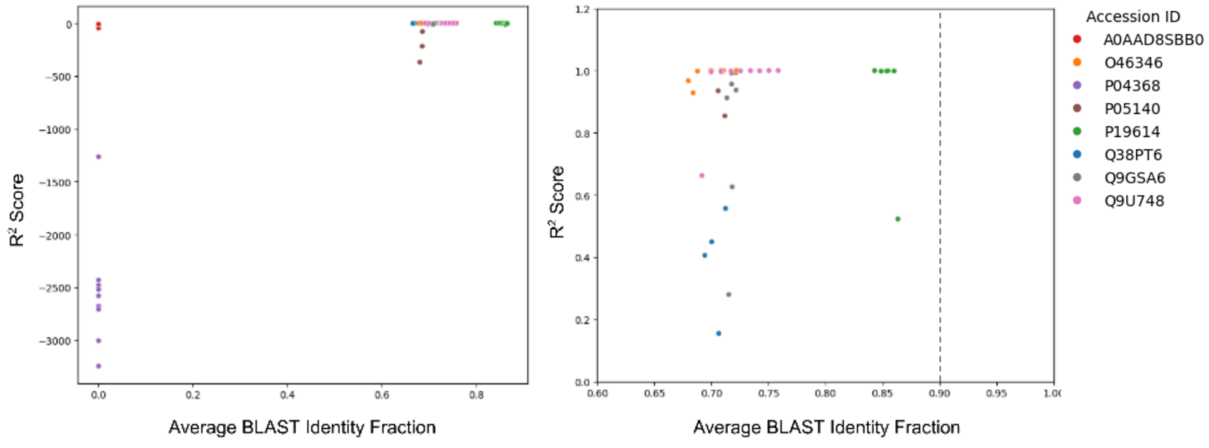


**Figure D. $R^2$ score vs protein identity fraction to activity seed proteins in training dataset.** Due to the potential similarity between anchor protein mutants and seed proteins, the training corpus was subset to limit the mutants based on their mutation percentage. Activity models were trained on each of the subsets, and the resultant ensemble R2 scores were shown for each seed protein. For each of the subsets, the BLAST alignments were also conducted between each mutant and its associated seed protein. The average BLAST identity fraction is then computed for each subset and used in the plots to depict the similarities between the subsets (used for model training) and seed proteins (used for model testing). The right plot is a zoomed-in version of the original plot on the left. As expected, the R2 scores increase as the average identity fraction increased. Notably, the models were able to achieve a high R2 score between 0.7 and 0.85 identity fraction. The dotted line references the 0.9 identity fraction, where at least 7 amino acids need to mutated (average of 11.5). This indicates an objective window for the similarity of training sequences to achieve an adequate accuracy for the activity model.

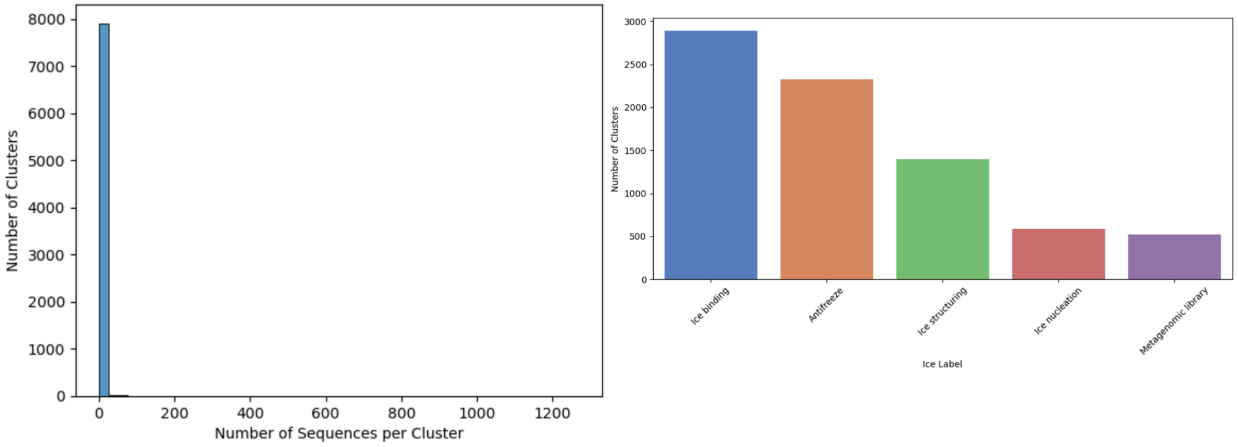


**Figure E. CD-Hit Analysis of Sequences for Ice Binding Training Corpus Similarity.** CD-Hit analysis was conducted at a 90% identity threshold for the ice binding protein and metagenomic library sequences to determine how similar the sequences are. Similar sequences are placed into clusters. The plot on the left depicts the distribution of cluster sizes. Of the 7959 clusters, a majority of them contain only 1 or 2 sequences, demonstrating that a great majority of sequences are unique. The plot on the right shows the most common ice labels associated with the clusters. Notably, there are over 2000 clusters with ‘Antifreeze’ as the most common label, significantly higher than the 400-500 protein sequences used by other models.

**Table 1. List of seed proteins and their targeted amino acids for activity model data augmentation.** Note that the targeted amino acids are used to mutate the associated anchor proteins to create mutants, which are then used for activity model training.

| Seed Protein Accession ID | Mutation-Targeted Amino Acid |
|---|---|
| P19614 | Alanine |
| P05140 | Alanine |
| Q9GSA6 | Alanine |
| Q38PT6 | Alanine |
| O46346 | Threonine |
| Q9U748 | Threonine |

**Table 2. List of metagenomic library seed proteins and their ice binding type (antifreeze protein (AFP) or ice nucleation protein (INP)).** Left: 1st generation seeds. Right: 2nd generation seeds.

| Accession ID | AFP/INP |
|---|---|
| A1YIY3 | AFP |
| A0A7U0TF31 | AFP |
| Q38PT6 | AFP |
| C7ED53 | AFP |
| P19614 | AFP |
| O16119 | AFP |
| P80961 | AFP |
| P05140 | AFP |
| E5LR38 | AFP |
| 6XNR | AFP |
| B2D1X8 | INP |
| O33479 | INP |

| Accession ID | AFP/INP |
|---|---|
| A1YIY3 | AFP |
| A0A7U0TF39 | AFP |
| Q38PT6 | AFP |
| UPI001643FB0F | AFP |
| P19614 | AFP |
| O16119 | AFP |
| Q9U747 | AFP |
| Q9GSA6 | AFP |
| E5LR38 | AFP |
| P04002 | AFP |
| B1P0S1 | AFP |
| B5T007 | AFP |
| A0A023J6X7 | AFP |
| Q086E4 | AFP |
| O46346 | AFP |
| B2D1X8 | INP |
| O33479 | INP |
